# Trial-by-trial fluctuations in decision criterion shape confidence

**DOI:** 10.1101/2025.09.04.674199

**Authors:** Robin Vloeberghs, Lara Navarrete Orejudo, Anne E Urai, Kobe Desender

## Abstract

Many of the choices we make are accompanied by a sense of confidence. Within classical Signal Detection Theory (SDT), confidence is conceptualized as the absolute distance between a decision variable and a decision criterion. The decision criterion is traditionally modelled as being stable over an experimental session. However, recent work challenges the notion of a static decision criterion, suggesting instead that the criterion undergoes trial-by-trial fluctuations. Combining SDT theory and model simulations, we predict that fluctuations in the decision criterion shape confidence. In 15 human decision-making datasets, trial-by-trial estimates of decision criterion were obtained with the Hierarchical Model for Fluctuations in Criterion (hMFC). Across all datasets, we confirmed our pre-registered hypothesis that confidence is shaped by single-trial criterion state. This effect was found in 14 out of 15 individual datasets, indicating a robust pattern across a variety of task paradigms and confidence reporting scales. Going beyond self-report, the shaping of confidence by criterion fluctuations was replicated in an implicit measure of confidence, RTs, and in two key neurophysiological markers, pupil-linked arousal and a neural signature of confidence. Our results demonstrate that variability in confidence, which has traditionally been treated as noise, actually reflects genuine sensitivity to the current state of the (fluctuating) decision criterion.

## Introduction

Every day we have to make a multitude of decisions. With each decision comes a subjective feeling of decision confidence, indicating the probability of having made the correct choice. Many computational models have helped to understand the mechanisms that give rise to people’s reported decision confidence when they make choices. Signal detection theory (SDT) is a classical and influential computational framework of decision making (Green & Swets, 1966), and easily extends to modelling decision confidence.

In its simplest form, SDT describes how observers classify noisy stimuli as belonging to one of two categories. The theory assumes that observers generate an internal representation of the relevant stimulus information, typically referred to as the decision variable (DV). To make a binary decision, this decision variable is compared against an internal decision criterion (Figure 1; static criterion, gray line). Because the brain is inherently noisy, repeated presentations of the same stimulus do not always result in the same internal representation. Instead, a distribution of decision variables is generated (Figure 1; two distributions, one for each stimulus). This inherent variability accounts for why participants can give different responses when presented with the same stimulus multiple times (Gold & Shadlen, 2007; Green & Swets, 1966; Macmillan & Creelman, 2005).

**Figure 1:**
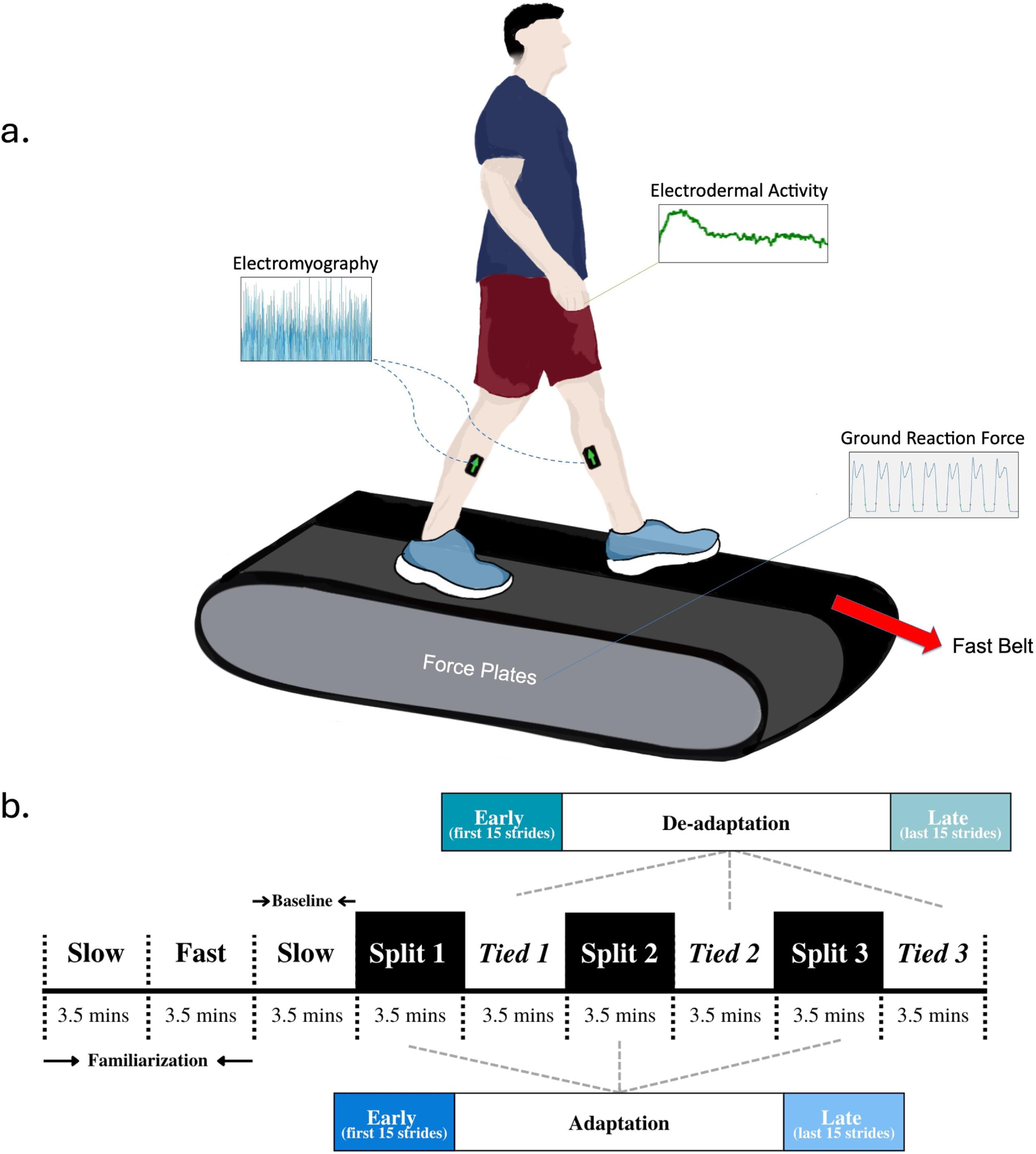
**A**. According to signal detection theory (SDT), choices are formed by comparing a decision variable to a criterion. An observer will respond “left” (squares) or “right” (circles), depending on whether the decision variable falls left or right to this criterion (grey line), respectively. Within SDT, confidence can be quantified as the absolute distance between the decision variable and the criterion (dotted lines), also known as the distance-to-criterion hypothesis. **B.** Whereas this criterion is typically assumed to be fixed across a series of trials (static criterion), here we investigate the consequences for decision confidence when this criterion slowly fluctuates across time. As illustrated, for identical decision variables, the predicted confidence can differ, depending on whether the criterion is static or fluctuating. **C.** Formal model simulations (6000 trials) from an SDT framework with a fluctuating criterion (i.e., as illustrated in Figure 1B), show a clear influence of criterion state on confidence. For left stimuli, confidence is higher when the criterion fluctuates towards the right. This pattern flips for right stimuli, with higher confidence when the criterion fluctuates towards left.

Going beyond binary decisions, SDT also explains how observers experience graded levels of confidence in the accuracy of their decisions. Specifically, decision confidence can be computed as the absolute distance between the decision variable and the decision criterion, with a larger distance indicating higher confidence (Hebart et al., 2016; Kepecs et al., 2008; Shekhar & Rahnev, 2021; Urai et al., 2017; Xue et al., 2024). This implementation is referred to as the distance-to-criterion hypothesis of confidence. Although this approach can be traced back already to the early proposals of SDT (cf. the rating experiment; Green & Swets, 1966) in recent years this approach has been popularized within the perceptual metacognition literature.

A key component in signal detection theory is the decision criterion to account for biases (i.e., when one response option is chosen more often than the other). For example, shifting the criterion to the right will result in overall more ‘left’ responses, while leaving the overall sensitivity unaffected. A large body of empirical work shows that human observers strategically shift the decision criterion in response to external feedback (Haddara & Rahnev, 2022), the base rate of stimuli (Sánchez-Fuenzalida et al., 2023), task instructions (Kloosterman et al., 2019), expectancy (Bang & Rahnev, 2017), previous choices (Treisman & Williams, 1984; Urai et al., 2019), and monetary rewards (Frithsen et al., 2018). To assess whether an observer uses a shifted (or biased) decision criterion, its position is calculated based on the hit and false alarm rate. Crucially, by doing so it is implicitly assumed that the criterion is fixed, and thus that this (potentially shifted) criterion is the same for all trials (Figure 1A; static criterion). However, increasing evidence suggests that computational variables, like the decision criterion, are not static but instead fluctuate at the level of individual trials (Figure 1B; fluctuating criterion) (Ashwood et al., 2022; Cowley et al., 2020; Gupta & Brody, 2022; Mochol et al., 2021; Roy et al., 2021; Urai, 2025; Vloeberghs et al., 2025).

In the signal detection theory framework, trial-by-trial criterion fluctuations should also affect the computation of confidence, given that confidence is thought to reflect the distance between the criterion and the decision variable (Figure 1B). More specifically, when the criterion fluctuates towards the right, on average the distance between a decision variable favoring the “left” stimulus and the criterion will increase, leading to higher confidence. On the contrary, when the criterion fluctuates towards the left the distance between a decision variable favoring the “left” stimulus and the criterion will decrease, leading to lower confidence. Indeed, exactly this pattern is observed when formally simulating data from an SDT observer with a fluctuating criterion (Figure 1C, see Method and Results for detailed description). These simulations also show that given the same stimulus, confidence can vary quite substantially. Thus, this suggests that variability in confidence reports, which have traditionally been treated as noise (Shekhar & Rahnev, 2021), might actually reflect genuine sensitivity of confidence judgments to the current state of the (fluctuating) decision criterion.

Here, we build on current developments in computational modeling which allow to effectively measure trial-to-trial fluctuations in the decision criterion. Vloeberghs and colleagues (2025) developed the hierarchical Model of Fluctuations in Criterion (hMFC), which uses a hierarchical Bayesian estimation approach to accurately recover criterion estimates at the single trial level. This new modelling framework now makes it possible to put the hypothesis to the test.

In sum, the current study aims to examine whether self-reported confidence ratings, reaction times, and neurophysiological measures of confidence are influenced by trial-to-trial fluctuations in the decision criterion. We first report results from a pre-registered analysis of 15 behavioral datasets, which confirmed the influence of criterion fluctuations on confidence ratings. Analyzing these datasets individually, the effect was found in 14 out of 15 datasets. In two additional datasets, these findings were replicated using an implicit measure of confidence, namely reaction times, and two neurophysiological markers of confidence, namely pupil size and a neural EEG signature of confidence. Together, these results suggest a robust effect that is present in a variety of paradigms and confidence scales and show that variability in both behavioral and neurophysiological measures of confidence is driven by fluctuations in the underlying latent decision criterion.

## Methods

### Open Science statement

A first exploratory analysis was performed on the data of Experiment 4 in Shekhar and Rahnev (2021) which is publicly available. Afterwards, we performed a pre-registered confirmatory analysis of 14 datasets from the Confidence Database (Rahnev et al., 2021). The preregistration can be found at https://osf.io/mxuv2. For ease of reading, in the remainder we will jointly discuss the results from the exploratory analysis together with the 14 confirmatory analyses. Finally, a pre-registered analysis regarding individual differences can be found in the Supplementary Materials. The analyses on the two additional datasets involving reaction times, pupil size, and the neural marker of confidence is not included in the pre-registration. All code and data is available at [insert link upon publication].

### Simulation study

To investigate how criterion fluctuations shape confidence, we simulated 6000 trials from an SDT observer. On each trial, a decision variable was sampled from a normal distribution with mean μ = [3, –2, –1, 1, 2, 3] and standard deviation σ = 1. The decision criterion varied across trials according to a first-order autoregressive process, AR(1), with autoregressive coefficient *a* = .9995 and noise standard deviation σ = .05. A binary response on each trial was obtained by comparing the decision variable to the decision criterion. Confidence was then quantified as the absolute distance between the decision variable and the decision criterion. To examine the relationship between confidence and criterion fluctuations, we used a linear regression model where confidence was predicted by a full-factorial model including as predictors the decision criterion, stimulus direction, and stimulus strength. Stimulus strength was defined as the absolute value of the mean μ of the normal distribution, while stimulus direction corresponded to its sign. For visualization purposes only, the decision criterion was divided into equally sized bins.

### Dataset Selection

As pre-registered, we only included studies that met the following inclusion criteria: 1) participants reported trial-by-trial confidence using a confidence rating scale, 2) each participant completed a minimum of 500 trials (i.e., to allow robust estimation of trial-by-trial criterion with hMFC), 3) the main manipulation involved the stimulus strength presented to participants on the screen, and 4) no trial-by-trial feedback was provided. Based on these criteria, a total of 14 datasets were selected (i.e., in addition to Shekhar & Rahnev (2021) which was used for the exploratory analysis). Note that in contrast to the pre-registration the datasets of Yeon et al. (experiment 1, unpublished) and Clark et al. (2018) are not included in the current analysis. The former was wrongly classified and only used one level of stimulus strength, and for the latter the fitting procedure of hMFC did not converge due the low number of subjects (n=4).

From the fifteen studies included, eight used Gabor patches as experimental stimuli, four studies used the random dot motion task (i.e., participants indicated whether they perceived left-or rightward movement within a cloud of moving dots), two studies asked participants to decide whether the left or right panel contained the most dots, and one study used ambiguous faces, which had to be classified as either looking fearful or happy. The majority of the studies, 10 out of 15, used a discrete 4-point confidence scale, whereas the other five studies either used a continuous scale or a 2-, 3-, or 6-point scale. Across studies, the number of subjects varied from 10 to 54, and the number of trials ranged between 504 and 3240. Thirteen subjects were excluded because they either performed at chance level or provided the same confidence rating on over 90% of trials. Six additional subjects were removed due to estimated criterion states larger than 10. Values this large saturate the sigmoid-transformed log-odds in hMFC, thereby jeopardizing the accuracy of the model estimates. In total, after excluding these subjects the final datasets comprised 379 subjects with 459.091 trials in total. A table with detailed information about each of the studies can be found in the Supplementary Materials.

To examine whether our findings extend to implicit measures of confidence and to neurophysiological correlates of confidence, we analyzed two additional datasets. For reaction time and pupil size, we used data from Urai et al. (2017), which included 27 human observers performing 500 trials of a two-interval forced choice (2-IFC) motion coherence discrimination task. In this task, participants judged whether a test random dot motion stimulus had higher or lower coherence than a reference stimulus, while their pupil sizes were recorded. Throughout the experiment, five stimulus strength levels were presented. Note that this dataset did not include confidence ratings. Reaction times were log-transformed and standardized. Pupil size was calculated as the mean baseline-corrected signal during the 250ms preceding feedback. More details on pupillometry preprocessing are provided in Urai et al. (2017). To assess the modulation of criterion fluctuations in a neural marker of confidence (error positivity), we reanalyzed data from Boldt & Yeung (2015). In this study, 16 participants decided which of two panels contained more dots and then rated their confidence on a six-point scale. Only a single stimulus strength level was used in this experiment. Single-trial error positivity amplitudes were calculated at Pz in a time window of 250ms to 350ms post-response, following Boldt & Yeung (2015). Technical details of the EEG preprocessing are available in the original study.

### Estimating the single-trial decision criterion using hMFC

The single-trial decision criterion was obtained by fitting the Hierarchical Model for Fluctuations in Criterion (hMFC) separately to each of the 15 datasets (Vloeberghs et al., 2025). This model uses a Bernoulli distribution to predict the binary response *y*_*t*_ at trial *t*:

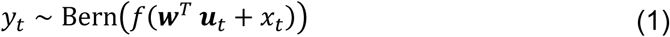

where the probability of a response is determined by a set of input variables *u*_*t*_, weighted by ***w***, and a latent variable *x*_*t*_ representing the decision criterion for that trial. The sigmoid transformation *f*(*Z*), restricts the probability between 0 and 1. The criterion state *x*_*t*_ is modelled as a first-order autoregressive model with intercept *b*, autoregressive coefficient *a*, and error variance σ^2^:

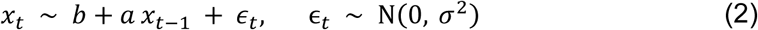

Adding the fluctuating value *x*_*t*_ to the Bernoulli log-odds is equivalent to comparing a decision variable to a fluctuating criterion. Note, however, that the estimated criterion *x*_*t*_, as outputted by hMFC, is a bias on the probability of making a certain response (e.g., *right* response). Thus, a positive criterion state will increase the probability of a *right* response, and therefore corresponds in SDT to a criterion shifted to the *left.* For ease of interpretation, in further analyses the sign of *x*_*t*_ is inverted such that its interpretation of *x*_*t*_ is in line with the traditional SDT interpretation.

In addition to the per-subject parameters (***w***, *b*, *a*, σ^2^), hMFC estimates hierarchical (group-level) distributions, allowing to share statistical strength across subjects to enhance the estimation accuracy of these individual parameters. Through this hierarchical structure, hMFC can accurately recover the model parameters, even with as few as 500 trials per subject. Most importantly, hMFC shows excellent recovery of the latent decision criterion trajectories (Figure 2A,B,C), providing accurate trial-to-trial estimates of the decision criterion. For full details on the model and its parameter recovery, we refer the reader to Vloeberghs et al. (2025).

**Figure 2:**
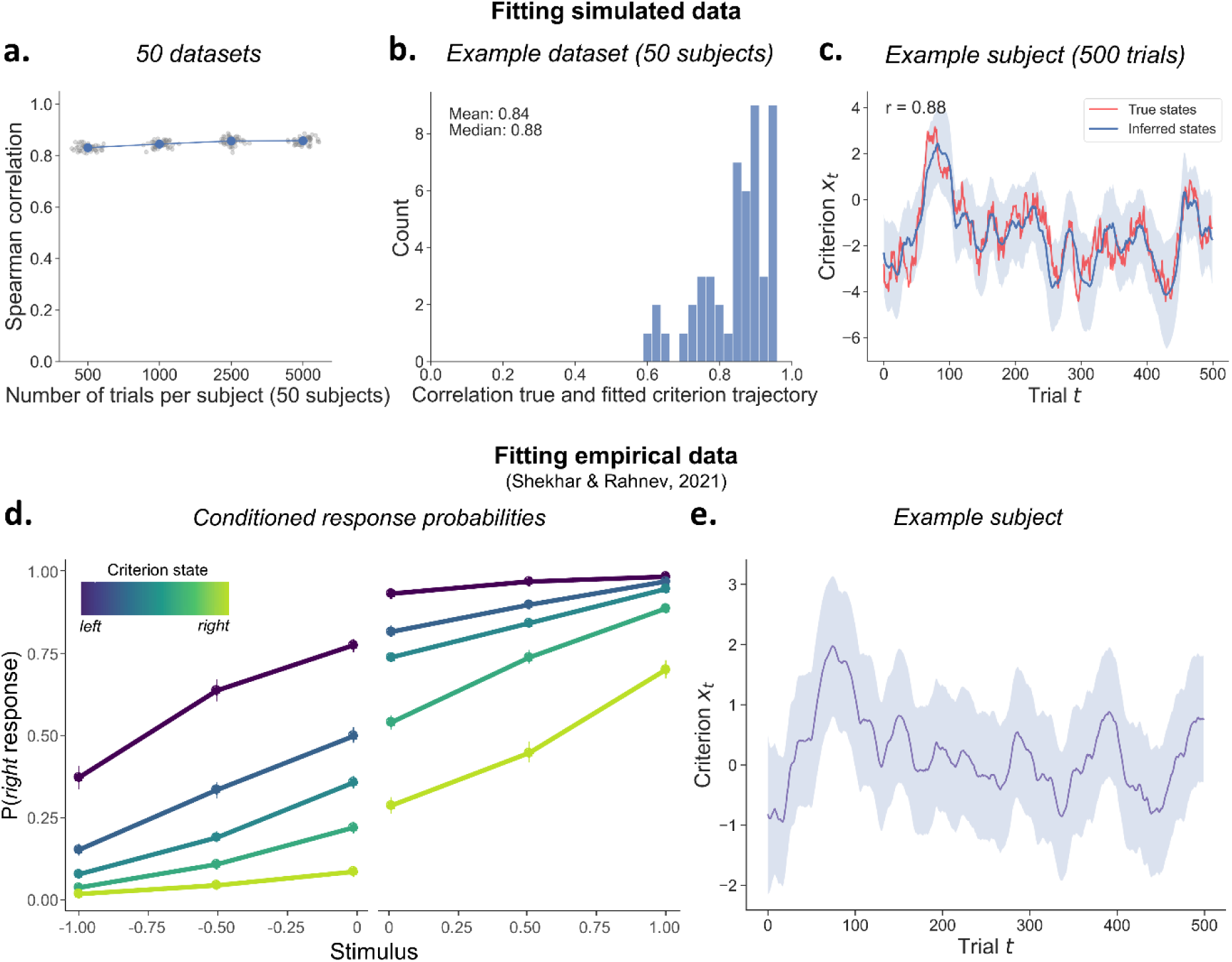
Estimating criterion fluctuations with hMFC on simulated and empirical data. **A.** The recoverability of the criterion fluctuations by hMFC was assessed in Vloeberghs et al. (2025) using simulated data with a varying number of trials per subject (500, 1000, 2500 or 5000 trials), with 50 subjects per dataset and 50 datasets per trial count. Each dot represents the correlation between the true and estimated criterion trajectory averaged over subjects in one dataset. The error bars are plotted but are very small. **B.** The correlation between true and estimated criterion trajectory for each subject (with 500 trials) in a representative simulated dataset. **C.** The true and estimated criterion fluctuations for a simulated subject with 500 trials. The shaded area indicates the 95% credible interval of the posterior at each trial. **D.** The empirical response probabilities of the data of Shekhar and Rahnev (2021, Experiment 4) conditioned on the binned trial-by-trial criterion. The more the criterion is shifted to the left, the more right responses are observed. **E.** The estimated criterion trajectory of an example subject (first 500 trials are shown) with the shaded area representing the 95% credible intervals.

In the current study, as input variables ***a***_*t*_ for hMFC we considered current trial stimulus, previous trial stimulus, and both response and confidence on the previous trial, together with its interaction. Note that in the data of Urai et al. (2017), used to investigate pupil size, self-reported confidence rating were absent and thus confidence on the previous trial could not be used as a predictor in hMFC. The model was run with four chains, each generating 5000 posterior samples with the first 500 samples discarded as burn-in. Convergence of the chains was assessed using the R-hat diagnostic. Fitting the full model with all the hierarchical (group-level) parameters resulted in convergence issues of the different chains (i.e., R-hat > 1.05) for the majority of the datasets. These issues were resolved by fixing μ_σ_2 and β_σ_2, the two parameters of the inverse gamma hierarchical prior for σ^2^, to the estimated values for Shekhar & Rahnev (2021), for which the model did properly converge. For consistency, all datasets were fitted with these two hyperparameters fixed.

### Predicting confidence with single-trial criterion state

To test the hypothesis that criterion fluctuations shape confidence ratings, we estimated three linear mixed effects models. The dependent variable in each of these models was reported confidence, which was rescaled to range between 0 to 1 for all datasets. In the first model, confidence was predicted based on the interaction between stimulus strength and stimulus direction. For example, in a random dot motion task, stimulus strength would correspond to the motion coherence and stimulus direction would refer to left-or rightward movement. Stimulus strength was recoded in each dataset to fall between 0 and 1, and stimulus direction was recoded as –1 and 1. In the second and third model, the single-trial criterion estimates, obtained with hMFC, were additionally included as a predictor variable. However, in the third model the estimated criterion trajectories were shuffled *across* subjects within each dataset (cf. session permutation method; Harris, 2020). Statistically comparing autocorrelated timeseries can introduce spurious significant effects when not sufficiently controlled for. To ensure that our models only picked up on confidence variability that are truly driven by trial-to-trial criterion fluctuations, we therefore employed the session permutation method by comparing Model 2 with Model 3. If confidence ratings are indeed influenced by criterion fluctuations, the second model including trial-by-trial criterion estimates should be favored over the other two models in a model selection procedure.

Model 1: Confidence ∼ stimulus strength * stimulus direction + (1 + stimulus strength | study / subject)

Model 2: Confidence ∼ stimulus strength * stimulus direction * criterion + (1 + stimulus strength + criterion | study / subject)

Model 3: Confidence ∼ stimulus strength * stimulus direction * shuffled criterion + (1 + stimulus strength + stimulus direction + shuffled criterion | study / subject)

First, we estimated the three models to all datasets *combined*. To account for the nesting of participants within studies when analyzing the datasets simultaneously, we used nested random effects. For model selection, it should be noted that the comparisons between Model 1 and Model 2, and between Model 2 and Model 3, were not conducted on the same dataset. Therefore, the results of these comparisons should not be compared directly. Specifically, the session permutation method used for Model 3 required all subjects within a dataset to have the same number of trials. To achieve this, within each dataset, we determined the cutoff based on the subject with the fewest trials and excluded any trials exceeding this count. As such, the estimated criterion trajectories could be shuffled *across* subjects. While the comparison between Model 2 and Model 3 was conducted on this truncated dataset, the comparison between Model 1 and Model 2 was performed on the full dataset. Second, we analyzed each dataset *separately* using Model 2, but without the random effects nested within each study.

To examine whether fluctuations in decision criterion influence reaction times and neurophysiological markers of confidence, we applied the same approach described above, replacing confidence ratings with reaction times, pupil size, or error positivity (Pe) amplitude as the dependent variable.

For all models, the random effects structure was obtained using an automated procedure which fits all possible random effect structures in parallel and selects the model with the lowest AIC. All statistical analyses were performed in R (R Core Team, 2023). The linear mixed effects models were fitted using the lme4 package (Bates et al., 2015).

## Results

### Model Simulations

To investigate whether and how confidence is shaped by criterion fluctuations we first simulated data from an SDT observer with the criterion varying from trial to trial (Figure 1C). As theoretically predicted, we find that confidence depends on the criterion. Crucially, the interaction between criterion and stimulus direction, which is central to our hypothesis, is significant (β = –0.423, SE = 0.035, *F*(1) = 3674.51, *p* < .001). In addition, we observe a significant three-way interaction between criterion, stimulus direction, and stimulus strength (β = –0.191, SE = 0.016, *F*(1) = 138.29, *p* < .001). While this three-way interaction is not crucial for our hypothesis, it suggests that the influence of criterion fluctuations on confidence itself depends on stimulus strength.

### Empirical Data

Following these simulations, we next investigated our hypothesis in empirical data, estimating criterion fluctuations using hMFC (Vloeberghs et al., 2025). hMFC can accurately recover the criterion fluctuations in simulated data, even with only 500 trials per subject (Figure 2A-C). In empirical data, the trial-by-trial criterion estimated by hMFC captures differences in response probabilities that are consistent with criterion shifts in SDT, as illustrated by the response probabilities conditioned on criterion state shown in Figure 2D. The more the criterion fluctuates to the right, the more left responses are observed. Conversely, a leftward shift in the criterion leads to more rightward responses. The model fits show stable and interpretable posterior distributions, with proper mixing of the chains of the Gibbs sampler (all R-hat < 1.05), reassuring that we sampled from the full posterior range (Suppl. Figure 1 and Suppl. Table 1). Supplementary Table 2 reports the mean, standard deviation, minimum, and maximum values, calculated per study, for the per-subject parameters of the autoregressive model that characterizes the criterion fluctuations. All together, these results ensure the validity of the hMFC estimates.

Next, we turn towards the key question of the current work: is confidence shaped by trial-to-trial fluctuations in decision criterion? In empirical data we observed the same confidence patterns as seen in the simulations (Figure 3A). Indeed, when inspecting Model 2, predicting confidence based on stimulus strength, stimulus direction, and criterion we found a highly significant interaction between stimulus direction and criterion, hereby confirming our main hypothesis (β = –0.043, SE = 0.002, *F*(1,456507) = 781.961, *p* < .001). The negative slope indicates lower confidence for a right stimulus (+1) when the criterion drifts towards the right (positive values), and higher confidence when the criterion drifts towards the left (negative values). The opposite is true for left stimuli (−1). Additionally, the three-way interaction was significant, similar to the simulations (β = –0.032, SE = 0.003, *F*(1,456276) = 123.330, *p* < .001). Although not the primary focus of the current study this shows that the interaction between stimulus direction and criterion was larger with stronger stimulus strength. Next, we found a significant effect of stimulus strength, with confidence increasing as stimulus strength increases (β = 0.587, SE = 0.114, *F*(1,14) = 26.715, *p* < .001). Lastly, the main effect of stimulus direction was borderline significant (β = –0.002, SE = 0.001, *F*(1,456483) = 3.665, *p* = .056). All other effects were not significant (all *p* ≥ .144). No multicollinearity problems were detected (all VIF ≤ 2.607).

**Figure 3:**
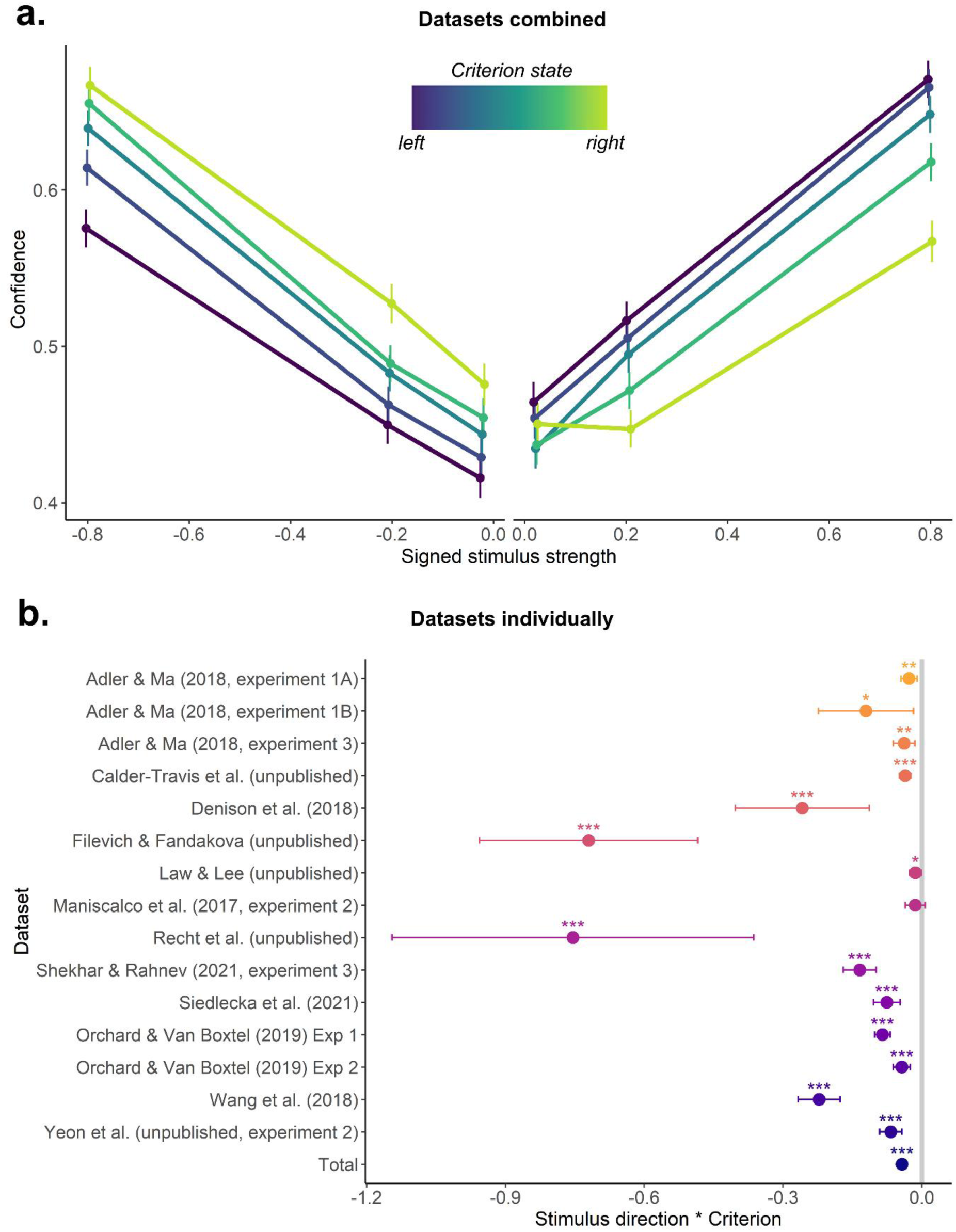
Trial-by-trial fluctuations in decision criterion shape confidence ratings in empirical data. **A.** As theoretically predicted, and in line with model simulations, decision confidence was modulated by trial-by-trial fluctuations in decision criterion. Specifically, when the stimulus direction was ‘right’ (positive signed stimulus strength) confidence increases as the decision criterion fluctuates towards the left (i.e. becomes more blue) and confidence decreases as the decision criterion fluctuates towards the right (i.e., become more yellow). Data are shown across the 15 datasets. Error bars reflect standard errors. For plotting purposes only, criterion states were binned and the average within each bin is shown. **B.** Parameter estimates for the crucial interaction between stimulus direction and decision criterion and their 95% confidence intervals. The estimates are obtained from models fitted separately to every dataset. The estimate plotted at the bottom, labelled “Total”, shows the parameter estimate from the model fitted to all datasets (taking into account the nesting of subjects in experiments). Note: * = p <.05, ** = p <.01, *** = p <.001.

To assess whether these findings reflect meaningful contributions of criterion fluctuations and not just statistical noise or model overfitting we turned to model comparison. Specifically, we tested whether a model incorporating trial-to-trial fluctuations in the decision criterion (Model 2) outperformed a simpler model that lacked these fluctuations (Model 1). Results provide unequivocal support that the more complex model including criterion is preferred (ΔAIC = –7482, ΔBIC = –7372, χ^2^(10) = 7502, *p* < .001). This indicates that a significant portion of variability in decision confidence can be explained by fluctuations in the decision criterion.

To further rule out the possibility that these results reflect spurious correlations that can occur between slowly drifting variables (Meijer, 2021), we implemented the session permutation method (Harris, 2020). In this approach, criterion fluctuations were randomly shuffled *across* participants (Model 3) and compared to the original Model 2. Once again, the comparison strongly favored Model 2 (ΔAIC = –8091, ΔBIC = –7848). In Model 3, the shuffled criterion fluctuations failed to account for any variability in confidence, as there was no interaction between stimulus direction and criterion (β = –0.002, SE = 0.002, *F*(1,137574) = 1.626, *p* = .202). Also the three-way interaction between stimulus direction, stimulus strength, and criterion was absent (β = 0.001, SE = 0.003, *F*(1,269873) = 0.182, *p* = .700). Only the main effect of stimulus strength remained significant (β = 0.592, SE = 0.116, *F*(1,14) = 26.228, *p* < .001).

In the previous, we have shown that *across* 15 datasets we found a highly robust association between confidence and the single-trial state of the decision criterion. Next, we further unraveled to what extent this pattern was present in each of the individual datasets. To this end, we fitted the model to each dataset separately (after omitting the random effects accounting for the nesting of subjects within studies). Importantly, the key two-way interaction between stimulus direction and criterion, central to our hypothesis, was significant in 14 out of 15 datasets (Figure 3B). The three-way interaction between stimulus direction, stimulus strength, and criterion, which also was predicted by the simulations, was significant in 10 out of 15 datasets, with 7 showing a negative slope and 3 showing a positive slope. In the remaining 5 datasets the interaction was absent (Suppl. Figure 2). In summary, although the three-way interaction is noisier and more difficult to detect at the dataset level, the consistent presence of the crucial interaction for our main hypothesis between stimulus direction and criterion in 14 out of 15 datasets demonstrates that the effect is robust and generalizes across diverse paradigms and confidence rating scales.

Having established that criterion fluctuations shape confidence ratings, we next explored whether this is also the case for an implicit measure of confidence (RTs) and for neurophysiological correlates of confidence. While ratings provide valuable subjective reports, they rely on participants’ introspection and may be influenced by biases or limitations in self-assessment. To complement this, we examined more objective indices of confidence that do not depend on introspective accuracy, namely reaction times, pupil size, and a neural marker of confidence (error positivity).

First, we consider reaction times. Reaction times are regarded as an implicit, non-self reported behavioral measure of confidence, with longer reaction times corresponding to lower confidence (Kiani et al., 2014; Urai et al., 2017). When predicting reaction times there was a significant effect of stimulus strength (β = –0.183, SE = 0.014, *F*(1,26.25) = 210.460, *p* < .001), with higher stimulus strength (i.e., easier stimuli) resulting in lower reaction times. Crucially, the interaction between stimulus direction and criterion was significant (β = 0.366, SE = 0.053, *F*(1,23.34) = 47.666, *p* < .001), indicating longer reaction times when the criterion fluctuates to the right for rightward stimuli (+1). Conversely, for leftward stimuli (−1), a rightward-shifted criterion resulted in shorter reaction times (Figure 4C). We further found a significant three-way interaction between stimulus direction, stimulus strength, and criterion (β = 0.026, SE = 0.007, *F*(1,1996.98) = 13.171, *p* < .001). All other effects were not significant (all *p* ≥ .156). It should be noted that this model showed some multicollinearity, with VIF scores reaching up to 10.25. Excluding the three-way interaction reduced VIFs to ≤ 3.185, while leaving all other results unchanged.

**Figure 4:**
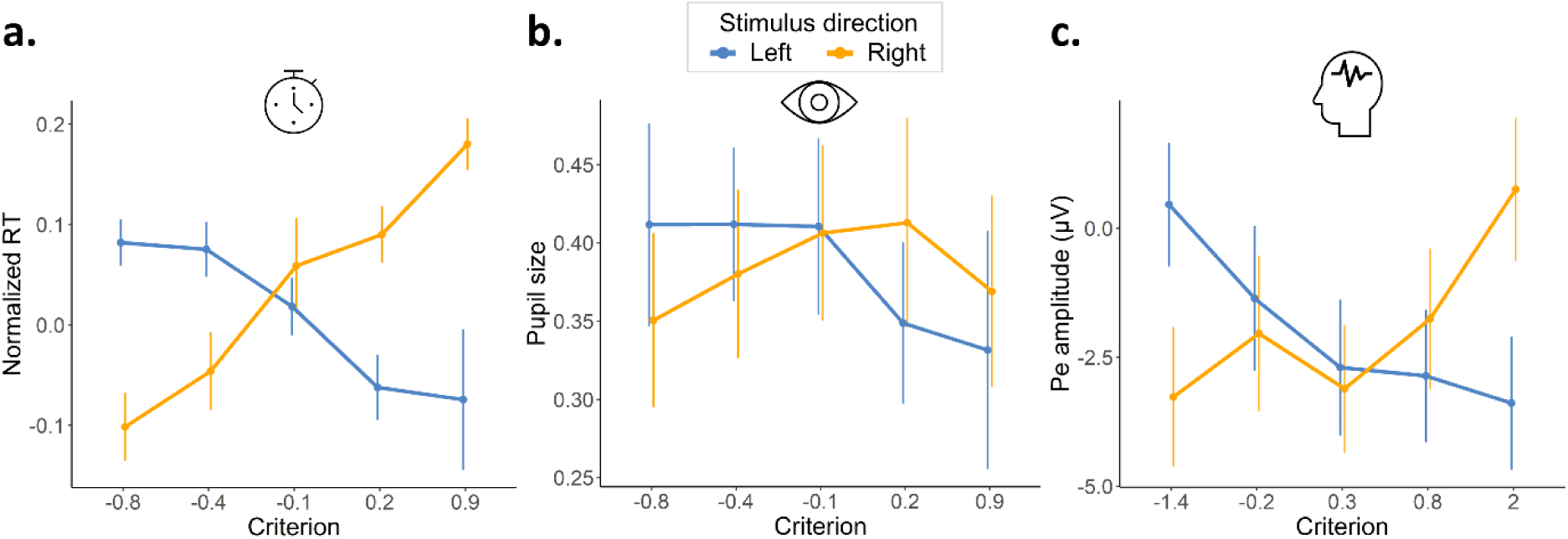
Criterion fluctuations shape neurophysiological measures of confidence. Pupil size (A), a neural marker of confidence (B), and reaction times (C) are all significantly predicted by the interaction between stimulus direction and criterion. Specifically, when the criterion fluctuates to the right (positive values) for a rightward stimulus (yellow line), longer reaction times, larger pupil sizes, and a higher Pe amplitude are observed, corresponding to low confidence. In contrast, when the criterion fluctuates to the left (negative values) for a rightward stimulus, lower values for all three measures are observed, indicating high confidence. Reaction times and pupil size show data from Urai et al. (2017), while Pe amplitude was taken from Boldt & Yeung (2015). In the latter dataset, only one level of stimulus strength was included, so it was not possible to show the data as a function of stimulus strength (as in Figure 3A). For consistency across panels, we therefore present only the two-way interaction. Error bars reflect the standard error over subjects. Note: * = p <.05, ** = p <.01, *** = p <.001.

Second, we examined a well-known neurophysiological marker of confidence: pupil size (Lempert et al., 2015; Urai et al., 2017). Phasic arousal, as quantified through pupil size, has been shown to be inversely related to confidence, with smaller pupil size reflecting higher confidence (Lempert et al., 2015). When predicting pupil size a significant main effect of stimulus strength emerged (β = –0.454, SE = 0.007, *F*(1,26) = 38.757, *p* < .001). Specifically, easier stimuli were associated with smaller pupil sizes, supporting the idea that pupil size is inversely related to confidence, since higher confidence is expected for easier stimuli. More importantly, we found that the crucial interaction between stimulus direction and criterion was significant (β = 0.064, SE = 0.021, *F*(1,67112) = 8.992, *p* = .003). A right-shifted criterion was associated with larger pupil sizes for rightward stimuli compared to leftward stimuli, and vice versa (Figure 4A). Finally, also the three-way interaction was significant (β = 0.018, SE = 0.008, *F*(1,67123) = 5.307, *p* = .021), as well as the interaction between stimulus direction and stimulus strength (β = 0.013, SE = 0.005, *F*(1,67126) = 7.292, *p* = .007). No other effects were present (all *p* ≥ .294) and multicollinearity issues were absent (all VIF ≤ 3.161).

Lastly, we investigated a neural marker of confidence, namely the error positivity measured with EEG (Pe; Boldt & Yeung, 2015). The error positivity is sensitive to graded levels of confidence, with its amplitude decreasing as confidence increases (Boldt & Yeung, 2015; Desender et al., 2019). When predicting single-trial Pe amplitudes, we found the critical interaction between stimulus direction and criterion (β = 1.912, SE = 0.203, *F*(1,764.5) = 89.23, *p* < .001), suggesting higher amplitudes for a rightward stimulus when the criterion fluctuates to the right, and lower amplitude when it fluctuates to the left (Figure 4B). Also the main effect of criterion was significant (β = –0.476, SE = 0.150, *F*(1,13086.3) = 19.786, *p* < .001). In this experiment there was only one level of stimulus strength, so the main effect of stimulus strength was omitted from the model. No multicollinearity issues were present (all VIF ≤ 1.942). Note that the same key finding in the pre-registered analysis on 15 datasets (i.e., interaction between stimulus direction and criterion) was also observed when analyzing the confidence ratings of this study, (β = –1.082, SE = 0.086, *F*(1,12.95) = 156.931, *p* < .001), with no other effects present (all *p* ≥ .564).

For all these three markers of confidence, the model including criterion outperformed the model without them (all ΔAIC ≤ –78, ΔBIC ≤ –41), and it was also preferred over the model with shuffled criterion trajectories (all ΔAIC ≤ –25, ΔBIC ≤ –61). Together, these results demonstrate that behavioral and neurophysiological markers of confidence are shaped by fluctuations in the decision criterion.

## Discussion

Within signal detection theory, decision confidence is traditionally conceptualized as the absolute distance between a decision variable and a criterion, which is known as the distance-to-criterion hypothesis. Crucially, in all previous work the criterion is assumed to be static. However, an increasing body of research suggests that computational parameters, such as the decision criterion, are dynamic and fluctuate over trials. Based on the distance-to-criterion hypothesis, we thus predict that fluctuations in decision criterion influence confidence ratings. Using the recently developed hierarchical Model for Fluctuations in Criterion (hMFC, Vloeberghs et al., 2025), which allows to obtain trial-by-trial estimates of the decision criterion, we tested this hypothesis. Across fifteen datasets, we found strong and consistent evidence for this hypothesis, demonstrating a robust effect independent of the specific paradigm or confidence scale. In addition, we replicated the effect in pupil size, in a neural marker of confidence and in RTs, suggesting that criterion fluctuations constitute a source of variability in confidence across both behavioral and neurophysiological measures.

The current work provides strong evidence in favor of the distance-to-criterion hypothesis of confidence: across 17 datasets we found that seemingly random variability in decision confidence actually tracks fluctuations in decision criterion. For confidence ratings, this effect was significant in 15 out of 16 datasets (including the dataset used for the neural marker of confidence), showing that this reflects a strong and highly replicable finding. Although the interaction was not significant in one dataset, the direction of the effect was consistent with the hypothesis. From a statistical point of view, finding one null effect across 16 datasets is not surprising: given experimental power of .8 it can be expected that 3 out of 16 studies show a null effect even when the effect is real. The one study were the effect was not found was not an obvious outlier in terms of design (i.e., a Gabor detection task both with 4-choice confidence reporting) nor estimated hMFC parameters (Table S3). Moreover, we replicated our findings across different neurophysiological measures of confidence indicating that the sensitivity to a fluctuating criterion reflects a fundamental aspect of how confidence is computed rather than merely a bias in how it is reported.

Our findings show that a substantial portion of variability in confidence ratings, which is typically interpreted as the results of some form of noise, can actually be attributed to fluctuations in the decision criterion. Another important consequence of quantifying confidence as the distance between the decision variable and a fluctuating criterion, is that confidence will naturally become autocorrelated across trials. This phenomenon has been described previously in empirical data, and is often referred to as the confidence leak (Rahnev et al., 2015). Based on the notion of fluctuations in the decision criterion, we are thus able using a fairly simple low-level explanation to explain both variability and autocorrelation in confidence ratings. Interestingly, in the literature these two observations (variability and autocorrelation) are usually explained via different mechanisms. Models not considering the possibility of fluctuations in the criterion attribute variability in confidence to some form of metacognitive noise that selectively perturbs the computation of confidence (De Martino et al., 2013; Jang et al., 2012; Mueller & Weidemann, 2008; Shekhar & Rahnev, 2021). To explain autocorrelation in confidence (i.e., confidence leak) Rahnev and colleagues (2015) proposed that participants dynamically update their confidence criterion on each trial depending on whether their confidence on the previous trial was higher or lower compared to average. Different from these two explanations, fluctuations in decision criterion provide a parsimonious account for these two key observations in the literature. Importantly, autocorrelation in decision criterion is not per se incompatible with the possibility of metacognitive noise or adaptations of confidence criteria based on previous-trial confidence. A key distinction, however, is that fluctuations in the decision criterion affect both responses and confidence ratings, whereas metacognitive noise and dynamic adjustments in confidence criteria only affect confidence ratings. Thus, future work could build on this distinction to further disentangle the respective contributions of fluctuations in criterion, metacognitive noise and dynamic adjustments to confidence criteria in explaining variability and autocorrelation in confidence.

In summary, our study provides robust evidence that confidence is modulated by trial-to-trial fluctuations in the decision criterion. This finding shows that variability in confidence judgments, which is often assumes to reflect noise, reflects genuine computation of confidence as distance-to-criterion. With this work new insights are provided into the computational underpinnings of decision confidence.

## Acknowledgement

We thank Annika Boldt for kindly providing us with the data of Boldt and Yeung (2015).

## Supplementary analyses

In a pre-registered exploratory analysis we investigated the relationship between confidence variability, measured as the standard deviation (SD) for each subject, the variability of the criterion fluctuations, and the hMFC parameters, *a* and σ^2^, which capture the temporal dynamics of the criterion fluctuations. All predictor variables were standardized.

Model 1: Confidence SD ∼ *a* * σ^2^ + (1 | study)

Model 2: Confidence SD ∼ criterion fluctuations SD + (1 | study)

In Model 1 the main effect of *a* was significant (β = –0.012, SE = 0.005, *F*(1,373) = 7.020, *p* = .008), suggesting that a higher value for *a* corresponds to lower variability in confidence. Both the effects of σ^2^ and the interaction between *a* and σ^2^ were not significant (all p ≥ .323). In Model 2 we found a significant main effect for the variability of the criterion fluctuations (β = – 0.008, SE = 0.003, *F*(1,368) = 5.788, *p* = .017). This indicates that subjects with larger variability in criterion showed less variability in their confidence ratings. No multicollinearity problems were detected (all VIF < 1.013). One possible explanation for these results is that subjects with high variability in criterion may be less engaged and attentive, and more often give the same confidence rating throughout the task. Note that including the per-subject criterion trajectory mean μ_*x*_, which is used in hMFC to estimate the intercept *b*, in Model 1 does not change the results. Therefore, we report the pre-registered analysis without μ_*x*_.

## Supplementary figures

**Supplementary Figure 1:**
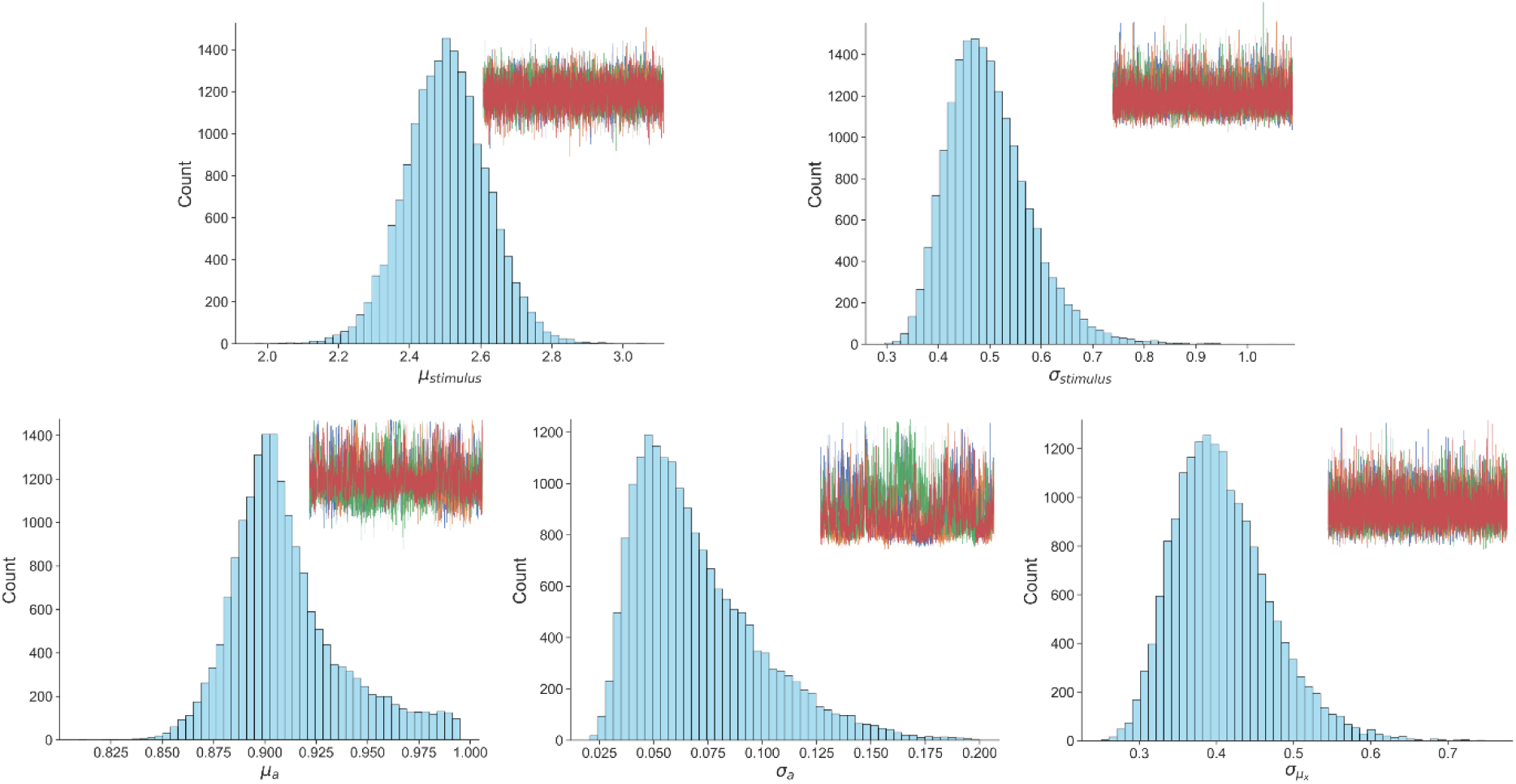
The posterior distributions of the hMFC hierarchical (group-level) parameters together with trace plots for Shekhar & Rahnev (2021). For each input variable *w* hMFC assumes a hierarchical (group-level) normal distribution with two estimated parameters: μ_*w*_ and σ_*w*_ . For the autoregressive coefficient *a* a truncated normal distribution constrained to [0,1] is specified with parameters μ_*a*_ and σ_*a*_. Lastly, the per-subject criterion trajectory mean μ_*x*_, which is estimated and allows a closed-form estimation for the intercept *b*, follows a zero-mean normal distribution with σ_μ*x*_ as standard deviation (see Vloeberghs et al. (2025) for more technical details). All parameters for each dataset have a R-hat lower than 1.05, which suggests a proper mixing of the chains of the Gibbs sampler. Note that for μ_*w*_ and σ_*w*_ the highest R-hat is shown out of the five input variables.

**Supplementary Figure 2:**
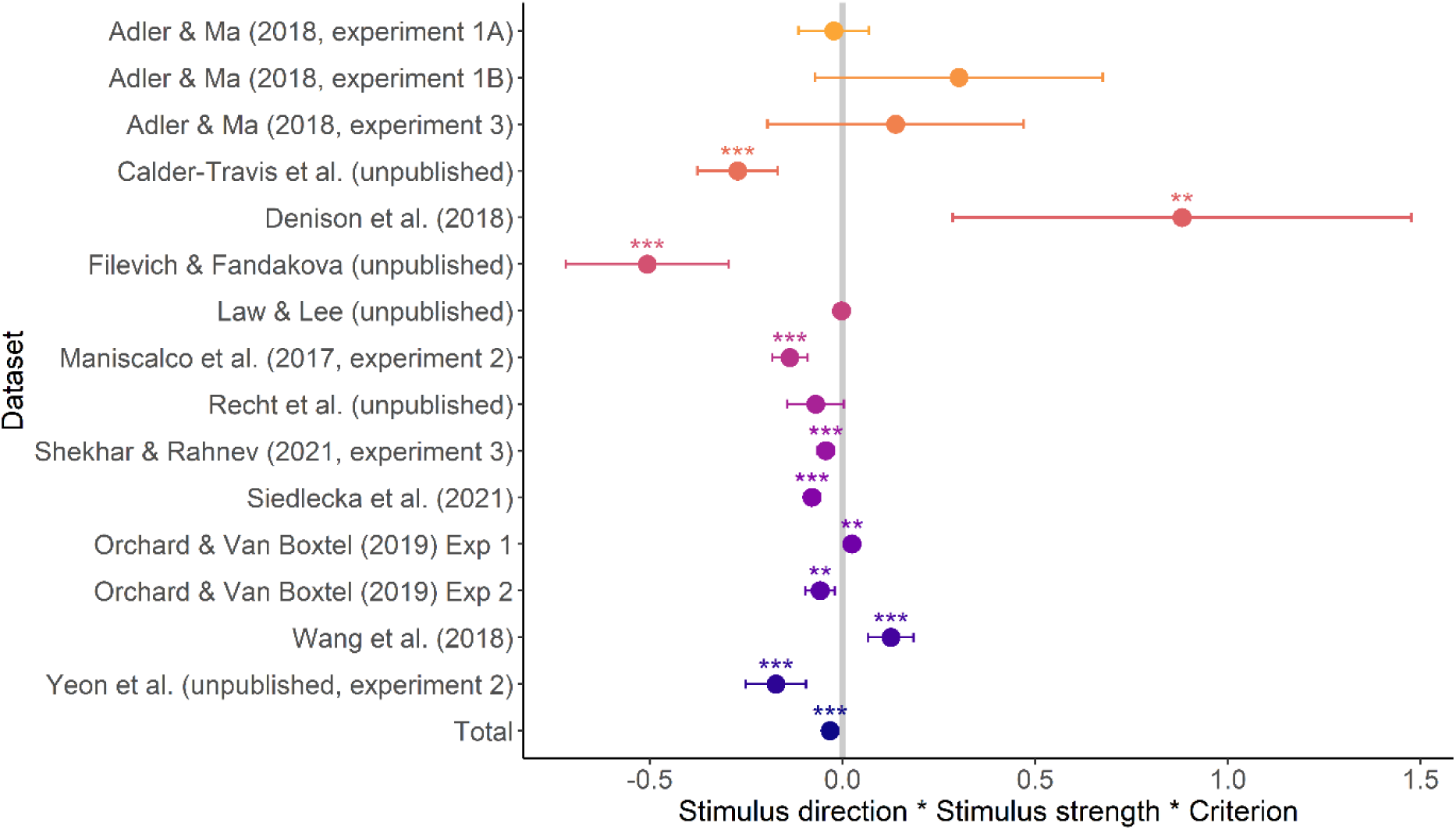
The three-way interaction of stimulus direction, stimulus strength, and criterion for each dataset separately. The parameter estimates and their 95% confidence intervals are shown obtained from models fitted separately to every dataset. The estimate plotted at the bottom, labelled “Total”, shows the parameter estimate from the model fitted to all datasets (taking into account the nesting of subjects in experiments). Note: * = p <.05, ** = p <.01, *** = p <.001.

## Supplementary tables

**Supplementary Table 1:** Overview of R-hat diagnostics per dataset to assess chain convergence of hMFC. All parameters for each dataset have a R-hat lower than 1.05, which suggests a proper mixing of the chains of the Gibbs sampler. Note that for μ_*w*_ and σ_*w*_ the highest R-hat is shown out of the five input variables.

| Study | $\mu_w$ | $\sigma_w$ | $\mu_a$ | $\sigma_a$ | $\sigma_{\mu_x}$ |
| --- | --- | --- | --- | --- | --- |
| Adler & Ma (2018, experiment 1A) | 1.001 | 1.000 | 1.035 | 1.026 | 1.000 |
| Adler & Ma (2018, experiment 1B) | 1.000 | 1.001 | 1.031 | 1.044 | 1.000 |
| Adler & Ma (2018, experiment 3) | 1.001 | 1.001 | 1.011 | 1.006 | 1.000 |
| Calder-Travis et al. (unpublished) | 1.002 | 1.004 | 1.008 | 1.018 | 1.000 |
| Denison et al. (2018) | 1.005 | 1.010 | 1.037 | 1.011 | 1.001 |
| Filevich & Fandakova (unpublished) | 1.001 | 1.001 | 1.008 | 1.008 | 1.000 |
| Law & Lee (unpublished) | 1.007 | 1.004 | 1.004 | 1.016 | 1.000 |
| Maniscalco et al. (2017, experiment 2) | 1.000 | 1.001 | 1.002 | 1.032 | 1.001 |
| Orchard & Van Boxtel (2019, experiment 1) | 1.008 | 1.024 | 1.020 | 1.008 | 1.001 |
| Orchard & Van Boxtel (2019, experiment 2) | 1.002 | 1.006 | 1.002 | 1.032 | 1.001 |
| Recht et al. (unpublished) | 1.003 | 1.001 | 1.047 | 1.009 | 1.000 |
| Shekhar & Rahnev (2021, experiment 3) | 1.003 | 1.001 | 1.006 | 1.039 | 1.000 |
| Siedlecka et al. (2021) | 1.001 | 1.000 | 1.001 | 1.005 | 1.000 |
| Wang (2018) | 1.002 | 1.001 | 1.019 | 1.000 | 1.001 |
| Yeon et al. (unpublished, experiment 2) | 1.007 | 1.011 | 1.003 | 1.002 | 1.000 |

**Supplementary Table 2:**
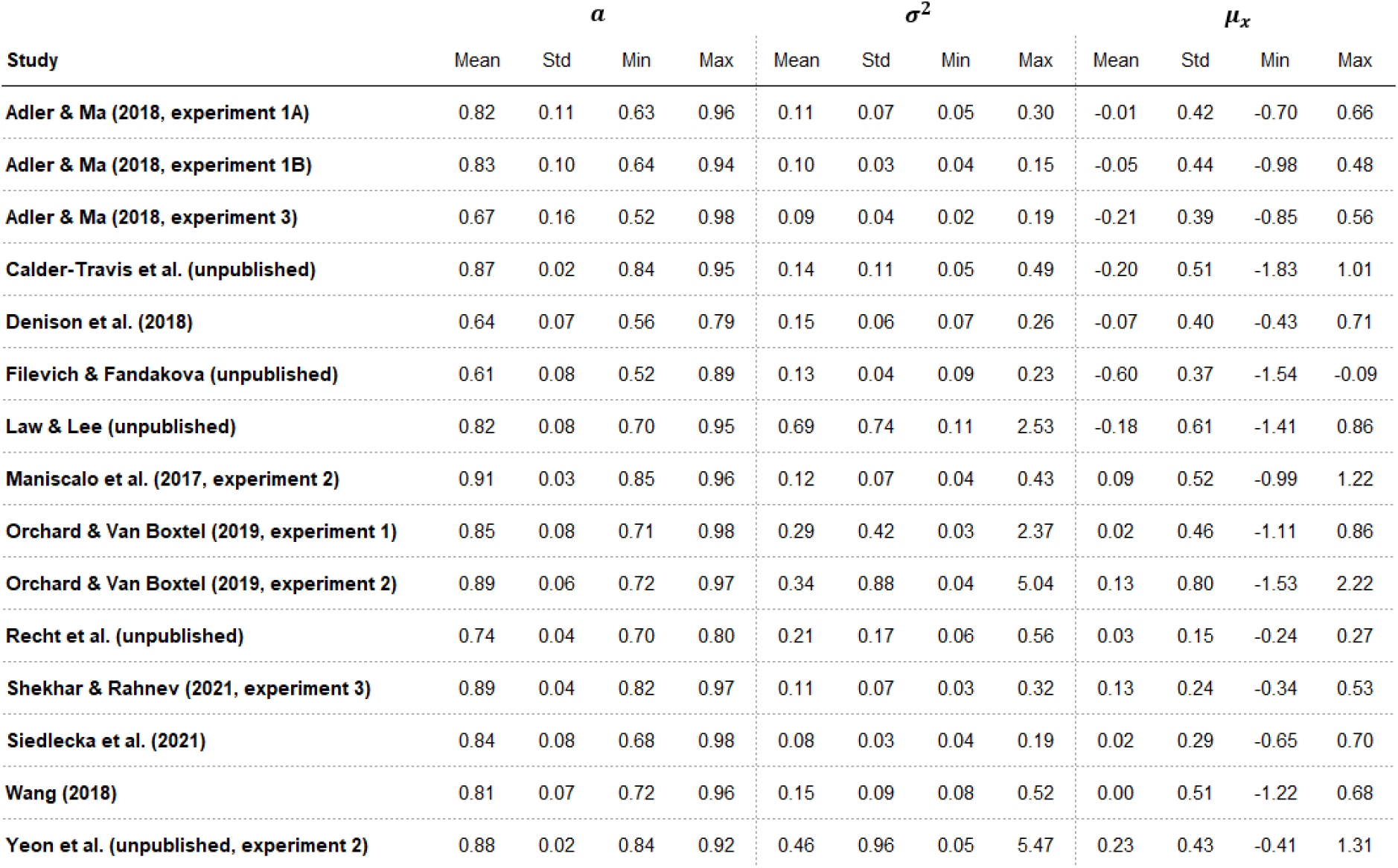
Overview per study of the mean, standard deviation, minimum, and maximum for *a*, σ^2^, and μ_*x*_. In hMFC the criterion fluctuations are assumed to follow a first-order autoregressive model. For each subject the autoregressive coefficient *a*, error variance σ^2^, and criterion trajectory mean μ_*x*_ is estimated. Note that μ_*x*_ is used for a closed-form estimation of the intercept *b* (for more technical details see Vloeberghs et al., 2025).

| Study | $\alpha$ | | | | $\sigma^2$ | | | | $\mu_x$ | | | |
| --- | --- | --- | --- | --- | --- | --- | --- | --- | --- | --- | --- | --- |
|  | Mean | Std | Min | Max | Mean | Std | Min | Max | Mean | Std | Min | Max |
| Adler & Ma (2018, experiment 1A) | 0.82 | 0.11 | 0.63 | 0.96 | 0.11 | 0.07 | 0.05 | 0.30 | -0.01 | 0.42 | -0.70 | 0.66 |
| Adler & Ma (2018, experiment 1B) | 0.83 | 0.10 | 0.64 | 0.94 | 0.10 | 0.03 | 0.04 | 0.15 | -0.05 | 0.44 | -0.98 | 0.48 |
| Adler & Ma (2018, experiment 3) | 0.67 | 0.16 | 0.52 | 0.98 | 0.09 | 0.04 | 0.02 | 0.19 | -0.21 | 0.39 | -0.85 | 0.56 |
| Calder-Travis et al. (unpublished) | 0.87 | 0.02 | 0.84 | 0.95 | 0.14 | 0.11 | 0.05 | 0.49 | -0.20 | 0.51 | -1.83 | 1.01 |
| Denison et al. (2018) | 0.64 | 0.07 | 0.56 | 0.79 | 0.15 | 0.06 | 0.07 | 0.26 | -0.07 | 0.40 | -0.43 | 0.71 |
| Filevich & Fandakova (unpublished) | 0.61 | 0.08 | 0.52 | 0.89 | 0.13 | 0.04 | 0.09 | 0.23 | -0.60 | 0.37 | -1.54 | -0.09 |
| Law & Lee (unpublished) | 0.82 | 0.08 | 0.70 | 0.95 | 0.69 | 0.74 | 0.11 | 2.53 | -0.18 | 0.61 | -1.41 | 0.86 |
| Maniscalco et al. (2017, experiment 2) | 0.91 | 0.03 | 0.85 | 0.96 | 0.12 | 0.07 | 0.04 | 0.43 | 0.09 | 0.52 | -0.99 | 1.22 |
| Orchard & Van Boxtel (2019, experiment 1) | 0.85 | 0.08 | 0.71 | 0.98 | 0.29 | 0.42 | 0.03 | 2.37 | 0.02 | 0.46 | -1.11 | 0.86 |
| Orchard & Van Boxtel (2019, experiment 2) | 0.89 | 0.06 | 0.72 | 0.97 | 0.34 | 0.88 | 0.04 | 5.04 | 0.13 | 0.80 | -1.53 | 2.22 |
| Recht et al. (unpublished) | 0.74 | 0.04 | 0.70 | 0.80 | 0.21 | 0.17 | 0.06 | 0.56 | 0.03 | 0.15 | -0.24 | 0.27 |
| Shekhar & Rahnev (2021, experiment 3) | 0.89 | 0.04 | 0.82 | 0.97 | 0.11 | 0.07 | 0.03 | 0.32 | 0.13 | 0.24 | -0.34 | 0.53 |
| Siedlecka et al. (2021) | 0.84 | 0.08 | 0.68 | 0.98 | 0.08 | 0.03 | 0.04 | 0.19 | 0.02 | 0.29 | -0.65 | 0.70 |
| Wang (2018) | 0.81 | 0.07 | 0.72 | 0.96 | 0.15 | 0.09 | 0.08 | 0.52 | 0.00 | 0.51 | -1.22 | 0.68 |
| Yeon et al. (unpublished, experiment 2) | 0.88 | 0.02 | 0.84 | 0.92 | 0.46 | 0.96 | 0.05 | 5.47 | 0.23 | 0.43 | -0.41 | 1.31 |

## Appendices

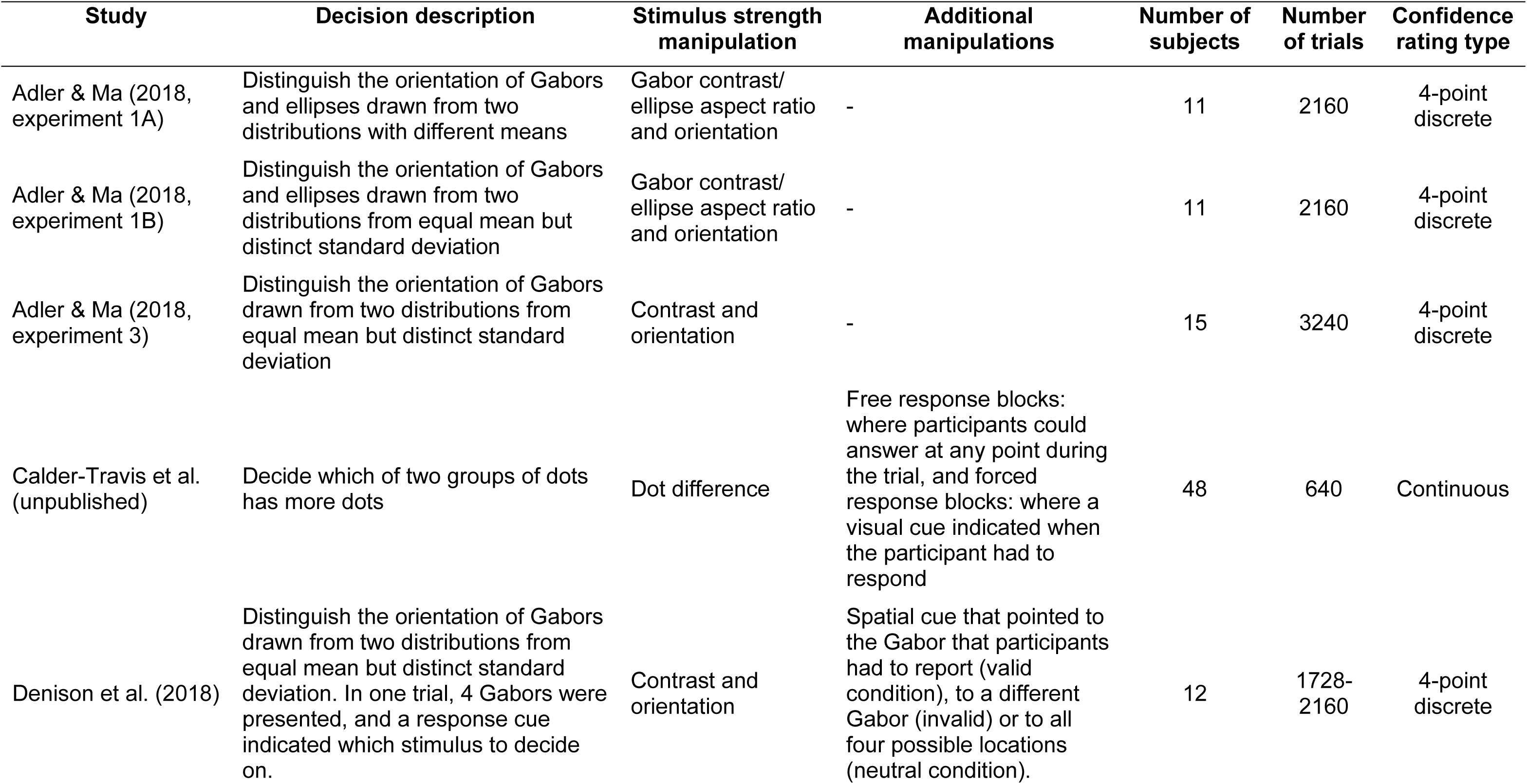
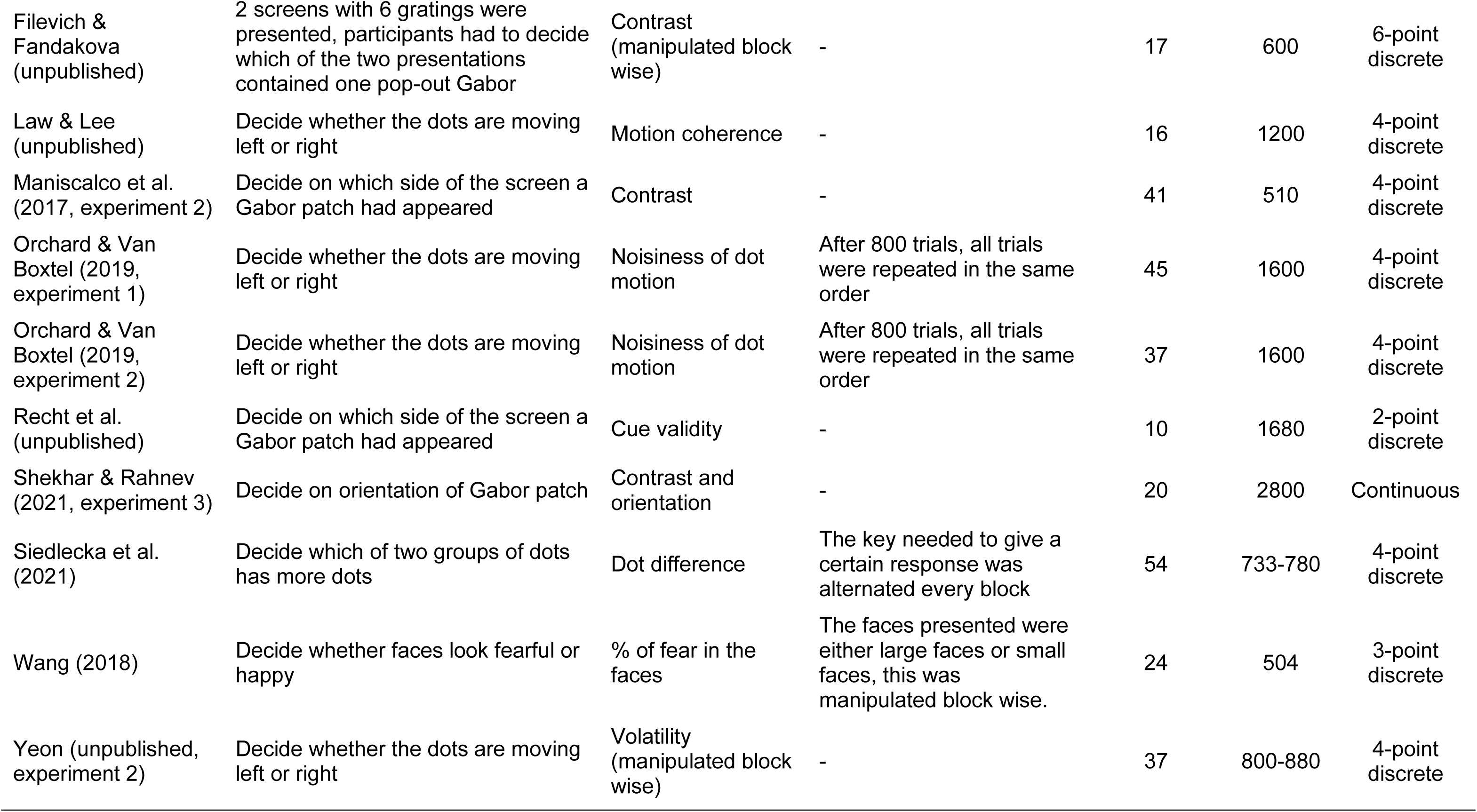

| Study | Decision description | Stimulus strength manipulation | Additional manipulations | Number of subjects | Number of trials | Confidence rating type |
| --- | --- | --- | --- | --- | --- | --- |
| Adler & Ma (2018, experiment 1A) | Distinguish the orientation of Gabors and ellipses drawn from two distributions with different means | Gabor contrast/ellipse aspect ratio and orientation | - | 11 | 2160 | 4-point discrete |
| Adler & Ma (2018, experiment 1B) | Distinguish the orientation of Gabors and ellipses drawn from two distributions from equal mean but distinct standard deviation | Gabor contrast/ellipse aspect ratio and orientation | - | 11 | 2160 | 4-point discrete |
| Adler & Ma (2018, experiment 3) | Distinguish the orientation of Gabors drawn from two distributions from equal mean but distinct standard deviation | Contrast and orientation | - | 15 | 3240 | 4-point discrete |
| Calder-Travis et al. (unpublished) | Decide which of two groups of dots has more dots | Dot difference | Free response blocks: where participants could answer at any point during the trial, and forced response blocks: where a visual cue indicated when the participant had to respond | 48 | 640 | Continuous |
| Denison et al. (2018) | Distinguish the orientation of Gabors drawn from two distributions from equal mean but distinct standard deviation. In one trial, 4 Gabors were presented, and a response cue indicated which stimulus to decide on. | Contrast and orientation | Spatial cue that pointed to the Gabor that participants had to report (valid condition), to a different Gabor (invalid) or to all four possible locations (neutral condition). | 12 | 1728-2160 | 4-point discrete |
| Filevich & Fandakova (unpublished) | 2 screens with 6 gratings were presented, participants had to decide which of the two presentations contained one pop-out Gabor | Contrast (manipulated block wise) | - | 17 | 600 | 6-point discrete |
| Law & Lee (unpublished) | Decide whether the dots are moving left or right | Motion coherence | - | 16 | 1200 | 4-point discrete |
| Maniscalco et al. (2017, experiment 2) | Decide on which side of the screen a Gabor patch had appeared | Contrast | - | 41 | 510 | 4-point discrete |
| Orchard & Van Boxtel (2019, experiment 1) | Decide whether the dots are moving left or right | Noisiness of dot motion | After 800 trials, all trials were repeated in the same order | 45 | 1600 | 4-point discrete |
| Orchard & Van Boxtel (2019, experiment 2) | Decide whether the dots are moving left or right | Noisiness of dot motion | After 800 trials, all trials were repeated in the same order | 37 | 1600 | 4-point discrete |
| Recht et al. (unpublished) | Decide on which side of the screen a Gabor patch had appeared | Cue validity | - | 10 | 1680 | 2-point discrete |
| Shekhar & Rahnev (2021, experiment 3) | Decide on orientation of Gabor patch | Contrast and orientation | - | 20 | 2800 | Continuous |
| Siedlecka et al. (2021) | Decide which of two groups of dots has more dots | Dot difference | The key needed to give a certain response was alternated every block | 54 | 733-780 | 4-point discrete |
| Wang (2018) | Decide whether faces look fearful or happy | % of fear in the faces | The faces presented were either large faces or small faces, this was manipulated block wise. | 24 | 504 | 3-point discrete |
| Yeon (unpublished, experiment 2) | Decide whether the dots are moving left or right | Volatility (manipulated block wise) | - | 37 | 800-880 | 4-point discrete |

